# Did succulents diversify in response to aridity? Evolutionary analyses of major succulent lineages around the world

**DOI:** 10.1101/2023.05.23.541957

**Authors:** Jamie Thompson, Santiago Ramírez-Barahona, Nicholas Priest, Tania Hernández-Hernández

## Abstract

The succulent syndrome is one of the most iconic life strategies in angiosperms, maximising water storage through a suite of adaptations to water-scarcity. Though succulence is considered a classic case of convergent evolution driven by shared environmental drivers, we lack a full understanding of whether the timing and drivers of the diversification of succulent lineages are, in fact, concordant. Here we analyse time-calibrated phylogenetic reconstructions of the seven richest lineages of succulents, and study diversification dynamics in relation to abiotic variables. Our analyses reveal different levels of synchronicity and relation with aridity. The impact of atmospheric CO_2_ on succulent macroevolution is varied. While transitions and radiations are especially concentrated in recent time, following a collapse of atmospheric CO_2_ ∼15 million years ago, CO_2_-dependent diversification is not supported in most lineages. With the exception of *Euphorbia*, we find that succulence elevates net diversification, though the effects on underlying speciation and extinction disagree. The phylogenetic distribution of transitions to succulence and rate shift increases suggest these phenomena are decoupled, indicating that succulence might not adhere to a classic key innovation model. We discuss that these evolutionary patterns reveal the ecological complexity of the succulent strategy, beyond simplistic interpretations as adaptations towards aridity.

## Introduction

As a life strategy, succulent plants have evolved a complex array of interrelated modifications, such as water storage tissue, shallow roots, a different type of photosynthesis known as Crassulacean Acid Metabolism (CAM), low surface area to volume ratio, and thick waxy cuticles amongst many other traits (Ogburn and Edwards, 2010; Arakaki et al, 2011; Griffiths and Males, 2017). Although hard to accurately delimit, the “succulent syndrome” is better described as a continuous spectrum, and confers high water use efficiency (WUE) compared to other life strategies (Males, 2017). The syndrome has been considered an iconic example of convergent evolution, having evolved repeatedly and independently across plant lineages (Arakaki et al, 2011; Griffiths and Males, 2017).

Of the approximately 290,000 angiosperm species, an estimated 3-5% or about 12,500 species are considered succulent, distributed across 690 genera in 83 families (Griffiths and Males, 2017). Some succulent clades show some of the highest diversification rates in the plant kingdom (Klak et al., 2004, Arakaki et al., 2011, Valente et al., 2013), and several of these hyperdiverse succulent clades are considered among the 30 core radiations by Magallón et al. (2019), being that they diversified rapidly into hundreds or thousands of species. These clades include the families Aizoaceae (>2,200 species), Crassulaceae (>1,500 species), Cactaceae (>1,500 species), the subfamilies Alooideae (>700 species) and Asclepiadoideae (>2,900 species), the tribe Agaveae (>300 species), and the megadiverse genus *Euphorbia* (>2,000 species) (Arakaki et al., 2011; Nyffeler and Eggli, 2010). Under this scenario, it seems straightforward to interpret the succulent syndrome as a key innovation allowing shifts to increased diversification rates in the lineages possessing it. However, succulence does not necessarily lead to increased rates of diversification, as exemplified by many succulent lineages that did not radiate dramatically into extant diversity, such as the monotypic family Halophytaceae (Anton et al., 2014) or the 13 species of Moringaceae (Olson and Carlquist, 2008).

Succulent angiosperms are found on all continents except Antarctica, including the Arctic tundra and above the Alpine snow line (Griffiths and Males, 2017), but geographic centres harbouring the highest succulent diversity exist in arid and semi-arid regions, most importantly Southern Africa and arid North America (Hernández-Hernández et al, 2014; 2023 *companying paper in review JXB*, Hilton-Taylor, 1996, Pillet et al., 2022). The elevated species richness of succulents in regions with limited water availability, and their remarkable climatic adaptations, have led aridification to take a central role in the hypotheses of the origin and diversification of these plants (Ehleringer and Monson, 1993; Arakaki et al., 2011; Edwards and Ogburn, 2012). Under these hypotheses, the exacerbation of environmental water stress during the aridification of terrestrial ecosystems would have provided succulent plants with an ecological and evolutionary advantage, permitting their higher diversification relative to non-succulents (Heyduk, 2022). Unfortunately, perhaps one of the hardest hypotheses to test is that of a relationship between aridification and the diversification of succulents, due to the absence of a fossil record for succulent plants, reliable paleoclimatic reconstructions of global aridification, and the lack of a timeframe for the evolution of succulence. For these reasons, a simple relationship between succulence and aridity has never been formally confirmed, especially at broad phylogenetic and spatial scales. Thus, questions about the timings of the origins of succulence and the importance of aridity and climate in their diversification remain unresolved (Grisebach 1872, Ellenberg, 1981; Von Willert et al; 1992).

The origin of the succulent CAM syndrome and diversification of succulent lineages has also been linked to atmospheric CO_2_ variation across geological time, due to the metabolic advantage of the improvement of net CO_2_ assimilation rate in environments of low atmospheric CO_2_ conferred by the CAM (Ehleringer and Monson, 1993; Arakaki et al., 2011; Hernández-Hernández et al. *in review JXB*).

Generally, the Cenozoic has witnessed a gradual decrease in atmospheric CO_2_ (Zhang et al., 2013), but exact variation in levels during this timeframe is heavily debated, with different proxies providing conflicting signals (Royer et al., 2001). A possible dramatic collapse commencing in the Middle Miocene at ∼15 Mya (Tripati et al., 2009) has previously been associated with radiations of cacti and other succulents, since both event timings seem to be coincident by visual inspections (Arakaki et al., 2011). Unfortunately, the link between atmospheric CO_2_ collapse and succulent radiations is still intuitive.

In this study, we further explore the evolutionary dynamics of major succulent lineages in relation to climatic variables related to arid environments and CO_2_ availability, which have been hypothesised as fundamental drivers of both succulent CAM plant and C4 grasses diversification (Arakaki et al. 2011; Spriggs et al, 2014; Bellasio et al, 2022). We use comprehensive phylogenetic frameworks to test multiple hypotheses regarding the tempo and mode of macroevolution in the seven richest succulent angiosperm clades. We first use a Bayesian method to detect diversification rate shifts and compare their temporal distribution with inferred timings of transitions to succulence. We then assess whether succulence promotes diversification by implementing sister-lineage comparisons for lineages with a single transition to succulence, and binary state-dependent speciation and extinction (BiSSE) models for lineages with multiple transitions (Maddison et al., 2007). We test for tree-wide dependence of continuously-varying diversification rates on CO_2_ under three scenarios of atmospheric change, accounting for the uncertainty associated around the hypothesised Miocene CO_2_-collapse (Beerling and Royer, 2011; Hansen et al., 2011). Finally, we estimated an aridity index for succulent species included in our study and applied quantitative state dependent speciation and extinction models (QuaSSE) to explicitly test for aridity-dependent diversification, and assess whether the shape of relationships differs among lineages (FitzJohn 2010). Although not definite, our comparative analytical framework improve our current understanding of the succulent CAM strategy and the complex set of ecological and evolutionary advantages it provides beyond simplistic one-size-fits-all explanations of plant survival and diversification under water scarcity.

## Materials and Methods

### Study groups, supermatrices and phylogenetic analyses

To provide a homogeneous framework to test diversification hypotheses, we started by reconstructing well-sampled dated phylogenies by maximising published nucleotide data, an approach commonly employed in comparative studies (Rabosky et al., 2018; Smith and Brown, 2018; Sun et al., 2020). We built supermatrices to construct robust time-calibrated phylogenies for each of the selected seven major lineages of succulent plants. Supermatrices were constructed from commonly sequenced loci used in plant phylogenetics and deposited in Genbank, with the exception of Asclepiadoideae and Cactaceae, for which we used recently published phylogenies made with the same supermatrix approach. For Asclepiadoideae, we pruned a recently reconstructed phylogeny of Apocynaceae (Fishbein et al., 2018), and for Cactaceae, we used a recent phylogeny constructed and time-calibrated with exactly the approach employed for the five remaining lineages included here (Thompson et al., 2023).

We sampled all available ingroup species in each lineage, plus multiple outgroups. Supermatrices were assembled with the OneTwoTree pipeline (Drori et al., 2018), which downloaded and clustered all available sequences with OrthoMCL (Li et al., 2003), and aligned clusters with Mafft (Katoh et al., 2002). Some clusters comprised fragmentary sequences with full-sequence counterparts, which we merged, keeping the full-sequence for species present in both clusters. In the cases where crown and stem divergence ages of the ingroup lineage were not available (see divergence time estimation below), we sampled outgroups more widely to capture nodes with available divergence estimates. Thus, for Agaveae we sampled more widely across Agavoideae, across Asphodelaceae for Alooideae, and across Euphorbiaceae for *Euphorbia*.

We extracted the alignments and inspected each of them visually for quality, trimmed poorly aligned regions using trimAl (--gappyout option) (Capella-Gutiérrez et al., 2009) and visually inspected alignments again. We concatenated alignments into supermatrices using AMAS (Borowiec 2016). Altogether, supermatrices comprised an average of 15.14 loci (ranging from 6 in Alooideae to 22 in Agaveae), with an average length of 15,096.30 bps (ranging from 4,997 in Alooideae to 28,197 in Asclepiadoideae; details of supermatrices as well as loci composition are available in Table S1, of the Supplementary Material.

### Scoring species as succulent

Similarly to CAM metabolism, succulence is best described as a continuous rather than a categorical trait, and plants can show a whole spectrum of succulence depending on how much water they are able to retain and in which tissue (Ogburn and Edwards, 2010; Griffiths and Males, 2017; Grace, 2019). For that reason, and to avoid ambiguity, we only considered as ‘succulent’ those species that have a significant amount of water storage tissue and have historically been considered as succulent, or in other words, show a dramatic unambiguous succulent syndrome. We followed a conservative approach and considered fleshy species or species showing mild succulence, or fleshiness, as non-succulent. For well-known genera and species, classification was straightforward (that is the case, for example, of succulent Cactaceae or *Agave*). In other cases, we resort to checklists, species descriptions in taxonomic literature, and visual inspection of photographs found in online biodiversity repositories (GBIF.org or Plants of the World). For Aizoaceae, we scored most Mesembryanthemoideae and Ruschioideae members as succulent, but scored most of the Aizooideae and Sesuvioideae as non-succulent. It is important to notice that although these were general trends, Mesembryanthemoideae includes some mild-succulent or fleshy members (e.g. *Mesembryanthemum cordifolium* or *M. crystallinum*). Similarly, the Aizooideae includes succulents (such as *Sesuvium portulacastrum*), and those were classified accordingly. For Asclepiadoideae and *Euphorbia*, we used published species-level assignments in Fishbein et al. (2018) and CITES succulents checklist (Carter et al., 2003) respectively, but we verified and edited the Asclepiadoideae classification based on genus-level data. We scored all members of Alooideae included in our phylogeny as succulent, as well as all members of Cactaceae with the exception of the leafy species of Pereskioideae (Anderson 2004). We scored members of Crassulaceae based on descriptions by Eggli (2003), only scoring as ‘succulent’ when succulence was explicitly described. A list of species included in our study and succulence classification are available on the Supplementary Materials.

### Estimation of divergence dates and transitions to succulence

Although complex statistical models for estimating topology and divergence times simultaneously were feasible for the smallest supermatrices (such as applying BEAST to Agaveae), the large size of most of them did not permit this due to computational power and times. Since the testing of hypotheses required the use of the same methodological approach and statistical framework for all lineages, we were restricted to using less computationally expensive methods. We reconstructed Maximum Likelihood (ML) phylogenies for each lineage and estimated divergence times using Penalized Likelihood (PL). To do this, we estimated the best partitioning strategy for each supermatrix using PartitionFinder2 under the greedy criterion (Lanfear et al., 2017) and used these to reconstruct ML phylogenies with RAxML v8.2.12 (Stamatakis, 2014) in the CIPRES gateway (Miller et al., 2011), assessing topological support by allowing rapid bootstrapping to halt automatically. Resulting best trees were used as input for divergence estimation with PL.

Since there is no fossil record for succulent or most arid adapted lineages, and to ensure congruence among the methodologies used for the estimation of divergence dates, we used secondary calibrations with divergence estimates obtained from Ramírez-Barahona et al. (2020). This is the most extensive estimation of dates available for angiosperms both in terms of the phylogenetic sampling and the fossil calibration strategy, with 238 fossil calibrations and a relaxed molecular clock approach. We extracted the dates in the relaxed complete set (Ramírez-Barahona et al, 2020), and use them to calibrate the seven trees with Penalized Likelihood (PL), with upper and lower bounds of higher posterior densities of divergence estimates. The crown and stem age of families was constrained, and for *Euphorbia*, the stem age of Euphorbioideae was used. Calibrations were implemented as uniform constraints in treePL (Smith and O’Meara, 2012). We identified the optimum priming strategy 100 times for each tree to find the most frequent cross-validation (cv) parameters. Then, we performed cv to find the smallest score, and reconstructed a calibrated phylogeny using the corresponding smooth value.

Following divergence estimation, we scored each species in the phylogenies as succulent or non-succulent, and used these data to reconstruct timings of transitions between states and study their diversification dynamics. To estimate the ages of transitions to succulence we used three approaches depending on the complexity of the lineage. For Alooideae, in which all species are succulent, we used the crown age as a conservative estimate. For Cactaceae, in which highly-succulent members form a monophyletic group comprising subfamilies Maihuenioideae, Opuntioideae and Cactoideae, we used the crown age of the succulent lineage. For the remaining lineages which have multiple transitions to succulence (Asclepiadoideae, Crassulaceae and *Euphorbia*), we estimated ancestral states using 100 SIMMAP simulations (Bollback 2006) in the R package phytools (Revell 2011), and visualised the distribution of transitions from non-succulent to succulent forms. We constrained the ancestral state of Agaveae to be non-succulent after an initial investigation in the wider Agavoideae tree from which we pruned Agaveae, because of the uncertainty encountered by analysing Agaveae alone. We replicated these plots for Alooideae and Cactaceae, but did not estimate the distribution of transitions to succulence for visualisation. Following this, we created state-dependent lineage through time (ltt) plots for all lineages, visualising against a randomly-selected ancestral state reconstruction.

### Characterising diversification-heterogeneity

We characterised diversification rate heterogeneity using Bayesian Analysis of Macroevolutionary Mixtures (BAMM, Rabosky 2014) after pruning outgroups. For each phylogeny, we estimated priors using the R package BAMMtools (Rabosky et al., 2014), and implemented a conservative prior of a single rate shift. To account for incomplete and imbalanced sampling, we specified sampling fractions either at the level of genus or subgenus (e.g. *Euphorbia*). We ran BAMM using four MCMC chains for 50 million generations, sampling chains every 5,000. We discarded a burnin of 10% and ensured convergence was met in every analysis (ESS of all parameters >200) using the R package coda (Plummer et al., 2006). Best shift configurations were estimated with BAMMtools, and the distribution of rate shift timings in the entire posterior sample of each clade was plotted as a distribution (n = 9,000). To assess if the rate shifts observed in our succulent lineages occurred with a distinctive pattern, we compared with rate shifts occurring in other non-succulent angiosperms. For that, we analysed the output from a BAMM analysis performed by Igea and Tanentzap (2020), where they used the megaphylogeny of >72,000 seed plants produced by Smith and Brown (2018). Here, incomplete sampling was accounted for with family-level sampling fractions and 1,000 samples of the posterior were saved. When analysing the BAMM output provided by Igea and Tanentzap (2020), we removed species of the lineages in this study, and plotted the distribution of rate shifts.

### Assessing atmospheric CO_2_ dependent diversification

We fitted time- and CO_2_-dependent Maximum Likelihood diversification models to the phylogenies after pruning outgroups, using the R package RPANDA (Morlon et al., 2016). For each phylogeny, six models were fitted where speciation rate varies exponentially with time or CO_2_, and extinction rate is zero, constant or varies exponentially with time or CO_2_. We did not fit constant models because their explanatory power is weak (Thompson et al., 2023) and we previously confirmed significant rate-heterogeneity in each lineage with BAMM. We also did not fit linear models because they are known to be problematic (Gamisch 2019; Morlon, personal communication). In all analyses we corrected for incomplete sampling by specifying the sampled fraction of known diversity.

We assessed evidence for three different impacts of atmospheric CO_2_, accounting for the uncertainty associated around the hypothesised Miocene CO_2_-collapse. We tested for tree-wide dependence of continuously-varying diversification rates on CO_2_ in each lineage with a robust continuous reconstruction which does not contain the Miocene collapse, favouring a more gradual decline (Hansen et al., 2013). We also tested for elevated tree-wide diversification ∼15 Mya in the absence of any paleoclimate reconstruction. Finally, we looked for biased distributions of timings of radiations within lineages, following ∼15 Mya. We used the high-resolution Cenozoic atmospheric CO_2_ curve reconstructed by Hansen et al. (2013). For Aizoaceae, which has an estimated crown age older than the maximum age in the CO_2_ reconstruction, we appended ∼12 Ma of the reconstruction by Bergman et al. (2004) to the reconstruction by Hansen et al. (2013). As a hypothesis test for a higher probability of CO_2_-dependent diversification models vs. time-dependent diversification models, we estimated evidence ratios (ER) using ΔAICc and Akaike weight (AICω), calculated as sum AICω CO_2_models / sum AICω time models (Thompson et al., 2023). Using a threshold ER of >20, we consider lineages with relative support for CO_2_-dependence of 20 as strongly supported. Conversely, lineages with ER <1/20 indicate strong support for time-dependence.

To test for a general tree-wide pattern of increased diversification after ∼15 Mya, we compared the fit of models of episodic and constant tree-wide diversification using the R package TESS (Höhna et al., 2015). We estimated marginal likelihoods for each model with stepping stone sampling, running 1,000 iterations with a burnin of 10% and using 50 stepping stones. We performed Bayes factor (BF) comparisons to select the best-supported model for each lineage. For lineages which supported the episodic model, we estimated parameters of this model for visualisation with 50 thousand MCMC generations, excluding a burnin of 10%.

### Testing for succulence-dependent diversification

We tested for state-dependent diversification under ML with BiSSE (Maddison et al., 2007) using diversitree (FitzJohn, 2012), in the lineages in which there are multiple transitions among non-succulent and succulent states. We compared four models differing by constraints: (1) one model in which parameters were estimated freely, (2) one in which speciation rates were constrained to be equal, (3) one in which extinction rates were constrained to be equal, and (4) one in in which speciation rates are equal and extinction rates are equal. For clades where the AIC score best supported any model of state-dependence, we visualised uncertainty by running BiSSE with 10,000 MCMC generations, with exponential priors, to estimate the posterior sample of state-dependent speciation rates. For lineages without multiple transitions to succulence (Alooideae and Cactaceae), we simply compared richness with their sister lineages.

### Aridity Index dependent diversification

Without a robust temporal reconstruction of the global rise of arid environments available, it has been difficult to accurately associate aridity with diversification. Our approach is to use modern aridity index values (mean annual precipitation/potential evapotranspiration) derived from spatial distribution data, to gain insights into the relationship between diversification rates and aridity. To model the effect of aridity index on diversification rate in succulent clades, we obtained median aridity index per species included in our phylogenies by using distribution data obtained from GBIF (GBIF.org), and aridity index climatic data obtained from CGIAR (Trabucco and Zomer, 2018). With the exception of Cactaceae, we downloaded available GBIF coordinates for species present in each tree and cleaned these with the R package CoordinateCleaner (Zizka et al., 2019). Records with location uncertainty beyond 10km, with zero coordinates and equal latitude and longitude were excluded. Additionally, records were removed where coordinates lay in seas, country capitals and centroids, and near biodiversity institutions. For Cactaceae, we used an existing dataset cleaned in a similar manner by Hernández-Hernández et al. (*companying paper in review JXB*).

We extracted Aridity Index (defined as mean annual precipitation/potential evapotranspiration) from CGIAR (Trabucco and Zomer, 2018) at a resolution of 30 arc-seconds, and estimated the median value for each species. We used QuaSSE to test the impact of median Aridity Index on diversification, implemented in diversitree (FitzJohn 2012). We ran seven analyses for each lineage and compared models with AIC scores to identify the best-supported model. Models in which character state had no impact on speciation rate were compared with models in which the effect was linear, sigmoidal, and modal. Additional linear, sigmoidal and modal models were estimated with the drift parameter, which estimates a directional trend. Incomplete sampling was accounted for with a sampling fraction, and aridity Index was either log transformed or log+1 in the case of Cactaceae, to improve normality.

As a simple general test of the hypothesis that succulents inhabit more arid regions than non-succulents, we estimated median aridity across angiosperms using a dataset of ∼15.8 million cleaned GBIF coordinates (Carta et al., 2021). We intersected this dataset with the megaphylogeny of ∼72,000 angiosperm species produced by Smith and Brown (2018), resulting in aridity index estimates for 59,999 species. For each of the seven succulent lineages, we calculated the median aridity index of all species. We replicated this for a randomly selected sample of number of similarly-sized lineages in the Smith and Brown megaphylogeny (all lineages with +/-10% of the species sampled in the focal succulent lineage), and performed a T-Test with the succulent-lineage median aridity index specified as the true value of the mean.

## Results

### Diversification rate heterogeneity and repeated evolution of succulence

By using a supermatrix approach we constructed robust time-calibrated phylogenies for each of the seven major groups of succulent plants to provide a well-resolved evolutionary framework. The fraction of species sampled per lineage in the supermatrices ranged from high (>69% for Cactaceae) to moderate (>21% in Asclepiadoideae and Aizoaceae). Although in some cases support values for phylogenies obtained were low (Figures S1-S7, Supplementary Material), all topologies are highly congruent with existing knowledge, including for the poorly-resolved Alooideae (Mort et al., 2001, Hernandez-Hernandez et al., 2014, Valente et al., 2013, Horn et al., 2014, Boucher et al., 2020, Jimenez-Barron et al., 2020). As expected, the extent of congruence of divergence estimates with previously available estimates varied among clades (Table S2, Suppl. Mat.).

We found strong evidence for rate-heterogeneity in diversification dynamics of all clades. Using BAMM, zero-shift models were not recovered in the posterior sample of any lineage, and the minimum number of shifts was one, recovered in Agaveae. Best rate shift configurations reveal all clades experienced multiple core shifts, ranging from two in Agaveae to 20 in Cactaceae. Rate shifts are distributed throughout each clade, e.g. in Cactaceae there are rate shifts in all major subfamilies (excluding the paraphyletic Pereskioideae) and in *Euphorbia* there are rate shifts in all subgenera. Tip-rates within clades varied remarkably, from 11.2-fold in Agaveae, to 70.3-fold in Asclepiadoideae (Figures 1a-g. Detailed BAMM output in Figures S8-14, Supp. Mat.).

**Figure 1.**
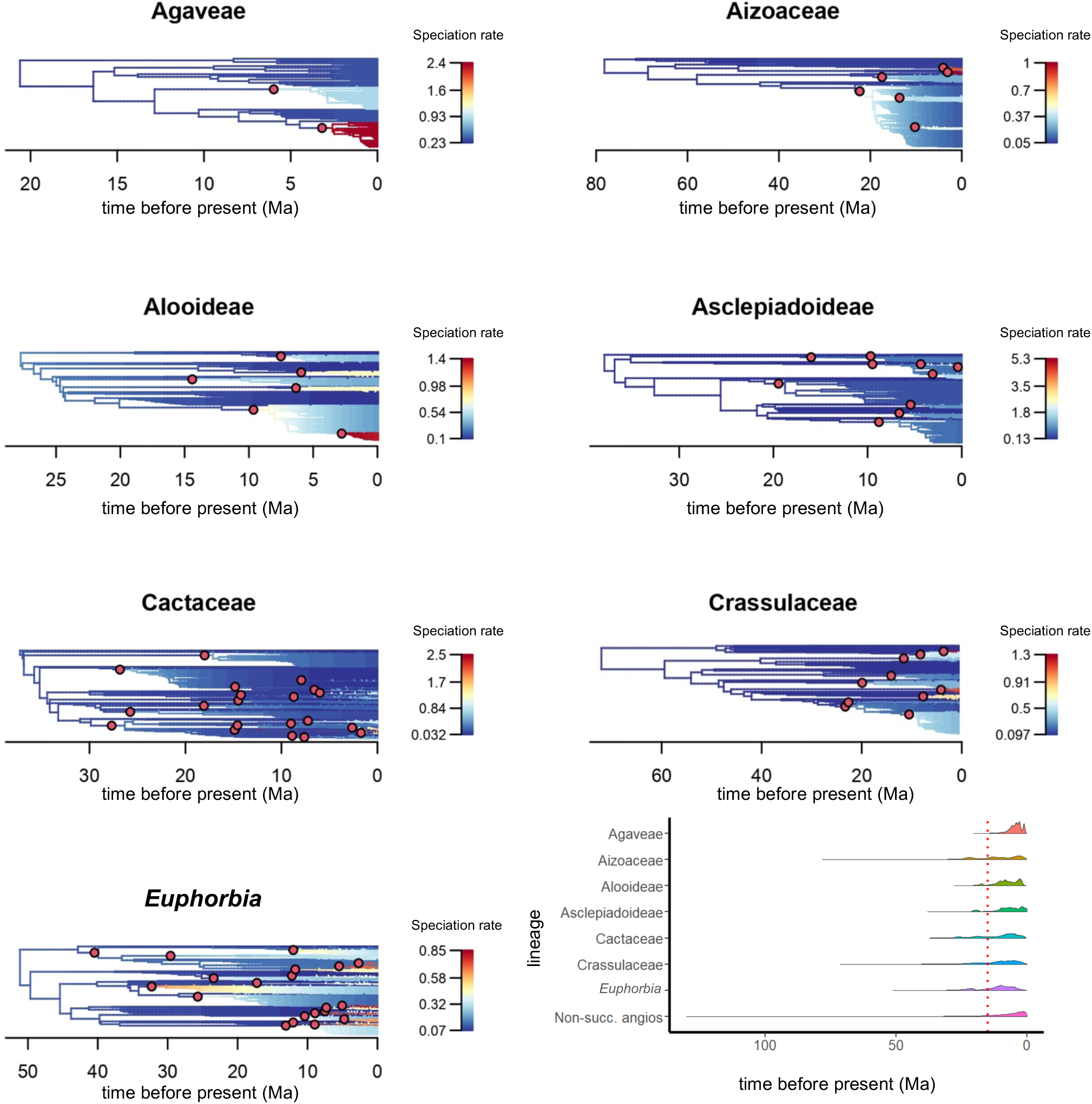
Heterogeneous diversification of the major succulent lineages. Diversification is shaped by multiple radiations which are especially concentrated following ∼15 Mya. The time-calibrated phylogeny is plotted for each lineage, and branches coloured by speciation rate estimated with BAMM (a-g). Rate shifts in the best shift configurations are plotted with red dots. Timings of the entire distribution of rate shifts in the posterior sample for each lineage are plotted as a half violin plot, alongside the posterior distribution of rate shifts detected in a BAMM analysis of ∼72,000 angiosperms (Igea and Tanentzap, 2020), with succulent lineages pruned (h). The red vertical line at 15 Mya denotes the hypothesised collapse of atmospheric CO_2_ (Tripati et al., 2009).

According to our reconstructions, the succulent syndrome emerged at different times in the seven lineages, though there is congruence among some of them (Figure 2). Succulence evolved no later than 36.86 Mya in Cactaceae, the crown age of succulent subfamilies, and 49.37 Mya in Alooideae, the crown age of the family (Figure 2, Table S2). In five clades (Agaveae, Aizoaceae, Asclepiadoideae, Crassulaceae and *Euphorbia*), there are multiple transitions into highly succulent lineages, permitting visualisation of uncertainty (Figure 2). Using 100 SIMMAP reconstructions we found an average of ∼4.26 transitions to succulence in Agaveae, ∼6.12 in Aizoaceae, ∼19.80 in Asclepiadoideae, ∼75.99 in Crassulaceae and ∼50.56 in *Euphorbia* (Table 1). Ancient transitions to succulence in these five lineages are reconstructed, but with low density. The highest density of transitions is found in recent geological time, after ∼20Mya, but especially after ∼10Mya (Figure 2h). Altogether, transitions to succulence are reconstructed throughout geological time, from the Paleocene in Crassulaceae to the Pleistocene in Asclepiadoideae.

**Figure 2.**
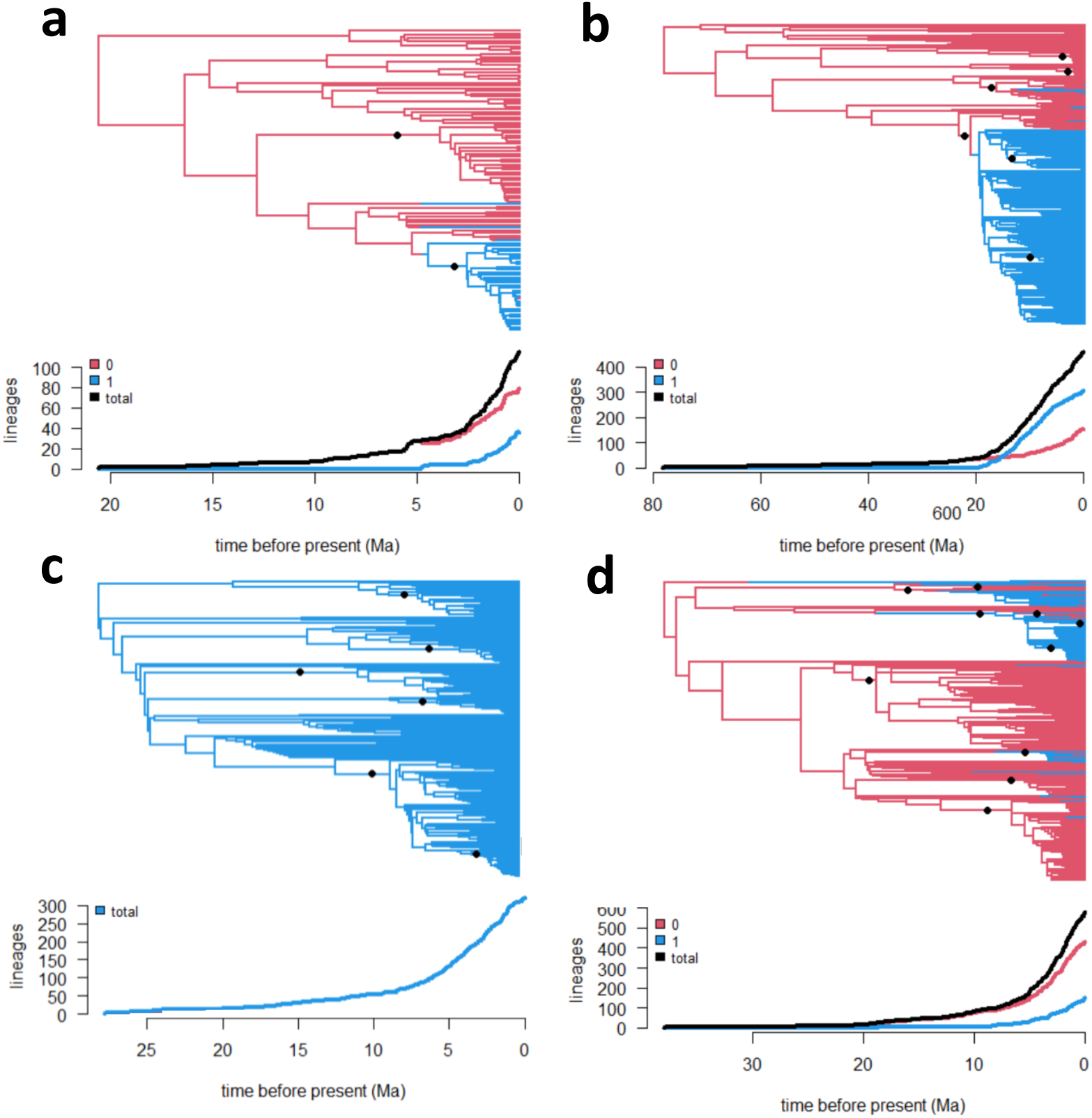
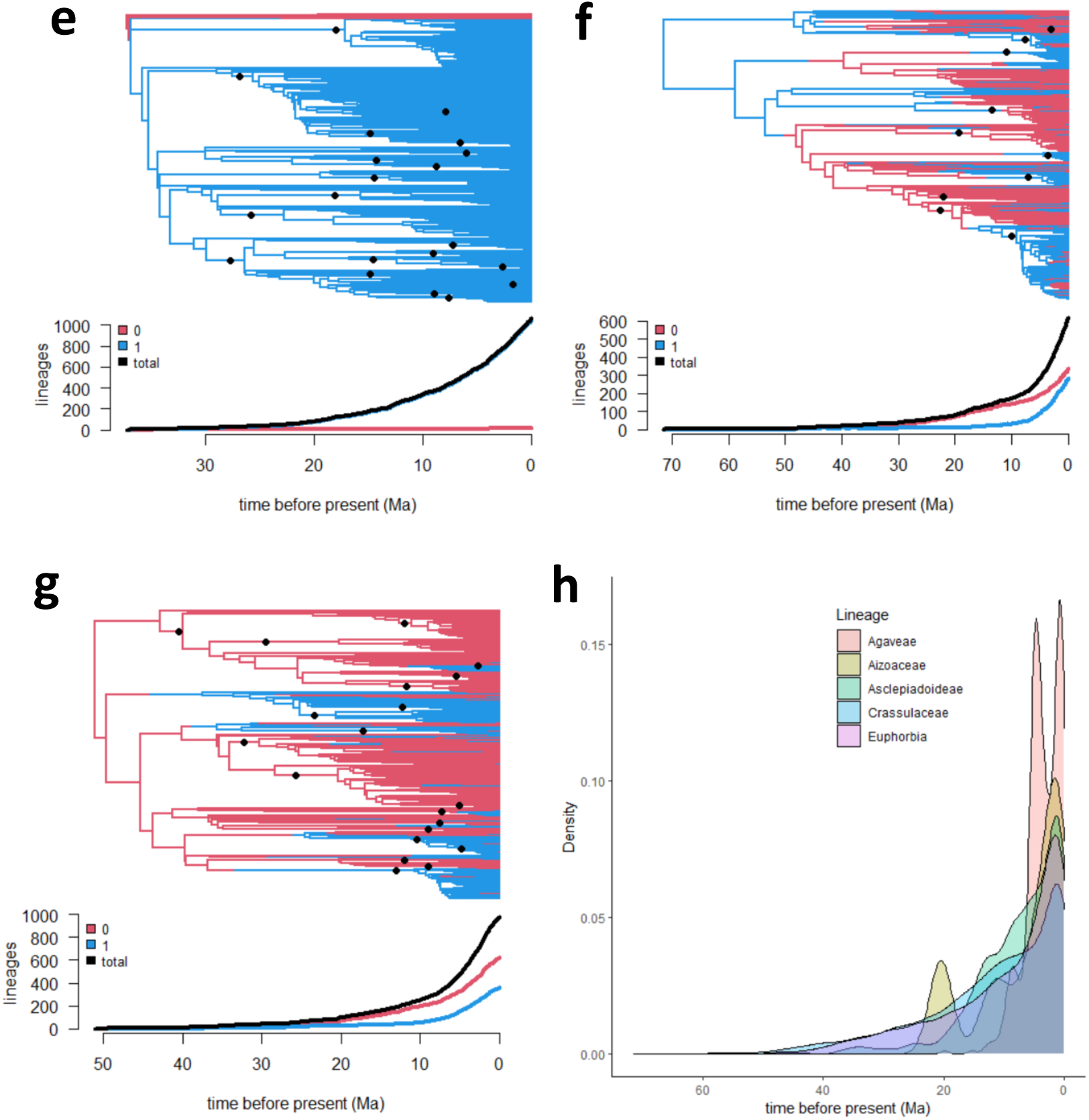
Evolutionary origins of succulence and macroevolutionary trajectories. Calibrated phylogenetic reconstructions showing the ancestral state reconstruction of succulence (blue lines), non-succulence (red lines) and state-dependent lineage through time (LTT) plots (below panels) for Agaveae (a), Aizoaceae (b), Alooideae (c), Asclepiadoideae (d). Black lines indicate overall LTT values. Cactaceae (e), Crassulaceae (f) and *Euphorbia* (g). The temporal density distribution of transitions to succulence in lineages with multiple transitions (h). A general decoupling between transitions to succulence and species radiations (black dots at internal nodes) is evident in most clades, as is rate-heterogeneity not associated with state-variation.

**Table 1:** Evidence for diversification associated to the succulent syndrome. For lineages shaped by a single origin of succulence (Alooideae and Cactaceae), a comparison of succulent richness against richness of their non-succulent sister lineage is presented. For lineages with multiple origins of succulence, the mean number of transitions estimated from 100 SIMMAP simulations is reported. The best-supported model is described, and the ΔAIC reported. For lineages supporting rates-different BiSSE models, the full parameters of Maximum Likelihood estimations are presented.

| Lineage | Number of transitions to succulence | Sister-lineage richness comparison | Best-supported BiSSE model and $\Delta AIC$ | Lambda0 (or shared lambda) | Lambda1 | Mu0 (or shared mu) | Mu1 |
| --- | --- | --- | --- | --- | --- | --- | --- |
| Agaveae | 4.26 |  | State-dependence, 8.42 | 0.282 | 0.754 | 0.078 | 0.050 |
| Aizoaceae | 6.12 |  | State-dependence, 29 | 0.522 | 0.220 | 0.484 | 0.000 |
| Alooideae | 1* | ~726, ~80 |  |  |  |  |  |
| Asclepiadoideae | 19.80 |  | State-dependence,<br>8.9 | 0.198 | 0.306 | 0.045 | 0.000 |
| Cactaceae | 1* | ~1,512, ~19 |  |  |  |  |  |
| Crassulaceae | 75.99 |  | State-dependence,<br>9.6 | 0.075 | 0.328 | 0.019 | 0.205 |
| *Euphorbia* | 50.56 | | No model<br>preferred<br>with $\Delta AIC >$<br>4, rates-<br>independent<br>model<br>reported | 0.285 | | 0.175 | |

### Decoupling between evolution of succulence and diversification rate shifts

Our analyses reveal that evolutionary transitions to succulence are often temporally and phylogenetically decoupled from diversification rate shifts in lineages. Diversification upticks either occurred more recently than major transitions to succulence (e.g. Agaveae, Figure 2a) or much earlier (e.g. Aizoaceae, Alooideae and Cactaceae, Figure 2b, c and e). Rate shifts are detected in the non-succulent sub-lineages, in every lineage possessing succulents and non-succulents except Cactaceae (Figure 1). Similarly, rate shifts are distributed within succulent sublineages, e.g. Aizoaceae, Alooideae, and Cactaceae (Figure 1b,c and e, respectively).

Increases in species richness seem to be associated with transitions to the succulent syndrome especially in those lineages with one major transition (Alooideae and Cactaceae), but this relationship is not evident in the lineages with multiple transitions (Table 1). In some cases, there is also no evidence that elevated diversification rates are coupled with succulence (e.g. Euphorbia, Figure 1-2). All Alooideae species are highly-succulent, and the richness (>700 sps) dwarves that of *Bulbine* (>80 sps), the non-succulent sister lineage (Chase et al., 2000). In Cactaceae, only the paraphyletic subfamily Pereskioideae is not highly succulent (Griffith, 2004), and the highly-succulent branch comprises over 99% of cacti richness (∼1,500 sps, Hernández-Hernández et al., 2014).

In the remaining five clades with repeated transitions between non-succulent and succulent forms, we used BiSSE to test for an association between succulence and diversification. In Agaveae, Aizoaceae, Asclepiadoideae and Crassulaceae, models in which speciation and extinction rates both differ by state are preferred (ΔAIC > 4, Table 1, Figure 3), but in *Euphorbia*, a rates-different model is not better supported (ΔAIC < 4, Table 1). In these lineages, net diversification is faster in succulents than non-succulents, but the impact on speciation and extinction rate differs. In Agaveae, Asclepiadoideae and Crassulaceae, speciation is faster in succulents than non-succulents, but the opposite pattern is found for Aizoaceae. In Agaveae, Aizoaceae and Asclepiadoideae, extinction rates are faster in non-succulents than succulents, but the opposite is found for Crassulaceae.

**Figure 3:**
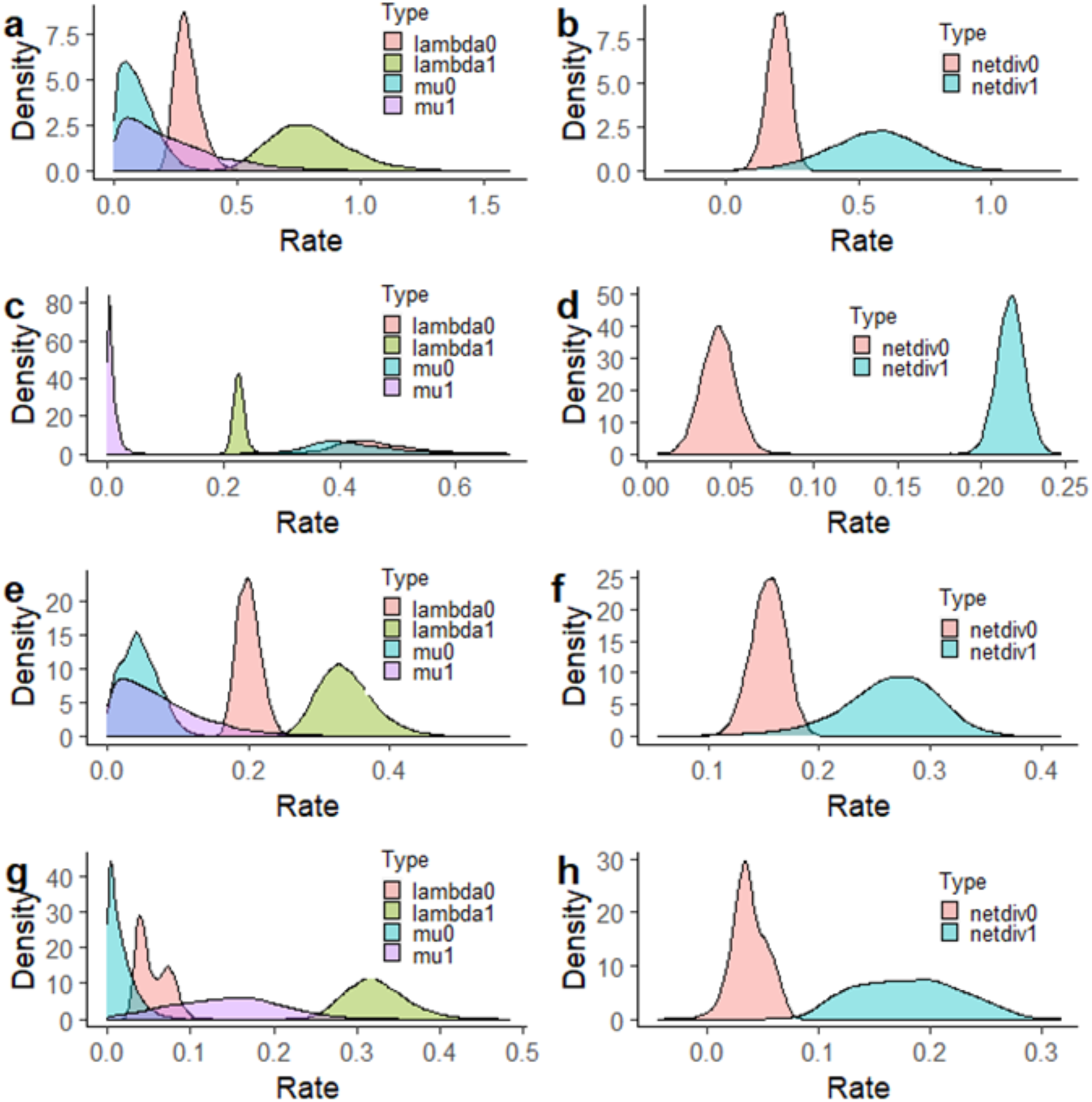
Elevated diversification of succulents shaped by differing speciation and extinction rates. The posterior distribution of BiSSE-estimated state-dependent speciation, extinction and net diversification rates for Agaveae (a, b), Aizoaceae (c,d), Asclepiadoideae (e,f), and Crassulaceae (g,h). Net-diversification rates are higher for succulents in these lineages, with speciation rates elevated by succulence in all lineages except Aizoaceae.

### Diversification rate shifts following the collapse of atmospheric CO_2_

We found little evidence for a general influence of CO_2_ on diversification in succulent plants. Different patterns of tree-wide CO_2_-dependent diversification are supported across the lineages for continuous temporal dependence (RPANDA), and discrete dependence (episodic model with a breakpoint at ∼15 Mya). RPANDA finds strong support for CO_2_-dependent diversification in Asclepiadoideae (ER 1704.59) and *Euphorbia* (ER 23.38), and for time-dependence in Aizoaceae (ER < 0.05). Neither time-nor-CO_2_ is strongly preferred for Agaveae, Alooideae, Cactaceae and Crassulaceae (ER 1.31, 0.43, 0.13 and 1.28, respectively). The best-fitting CO_2_-dependent model for Asclepiadoideae is one in which speciation rates vary exponentially with CO_2_, with no extinction, and for *Euphorbia* is one in which both speciation and extinction rates vary exponentially with CO_2_. An episodic diversification with a breakpoint at 15 Mya was strongly supported in Asclepiadoideae (BF 4.00) and substantially in *Euphorbia* (BF 2.97), although the nature of the relationship differed. For Asclepiadoideae, diversification rates are higher after 15 Mya (mean before 15 Mya: 0.07, mean after: 0.18) but for Euphorbia they are lower (before: 0.12, after: 0.11). A constant-rate model was moderately preferred for Alooideae (BF 4.39), and weakly preferred for Agaveae (BF 1.29), Aizoaceae (BF 1.12), and Cactaceae (BF 1.61). Neither model was preferred for Crassulaceae (BF 0.30).

Despite the lack of evidence for lineage-wide influences of CO_2_, we find a preponderance of diversification rate shifts following the hypothesised collapse of atmospheric CO_2_ at ∼15 Mya. While we found evidence of rate shifts that predate the global collapse of atmospheric CO_2_ at c.15Ma (Tripati et al., 2009), the majority are concentrated afterwards (Figure 1). The earliest rate shifts are recovered in the Eocene (56.0-33.9Mya), in Crassulaceae and *Euphorbia*. Oligocene rate shifts are recovered in Aizoaceae, Cactaceae, Crassulaceae and *Euphorbia* (c.33.9-23.0Mya). Miocene rate shifts are recovered in all clades (c.23.0-5.3Mya). Pliocene (c.5.3-2.6Mya) rate shifts are recovered in all clades except Agaveae, while only Asclepiadoideae and Cactaceae have rate shifts recovered in the Pleistocene (2.6Ma-11.7Kya). In total, there are 76 rate shifts recovered in the best shift configurations, with over two thirds occurring in the Miocene (51 rate shifts, 67%). Though we found an enrichment of rate shift subsequent to the collapse of atmospheric CO_2_, the rate shifts were not contemporaneous and occurred at levels similar to non-succulent plants (Figure 1h).

### Aridity correlates with diversification rates in different ways

Aridity-dependent diversification is supported in all seven clades, but the best-fitting models indicate different relationships (Figure 4, Table 2). Although QuaSSE cannot be considered to exactly explain the true relationship between continuous variables and diversification, it can estimate general trends (FitzJohn 2011). Sigmoidal relationships were found for Agaveae, Asclepiadoideae and Cactaceae, but the direction of this trend differs. In Agaveae, speciation rate is highest in lineages experiencing higher aridity and lower above a threshold aridity index of ∼0.33, while in Asclepiadoideae and Cactaceae, speciation rates are highest below the threshold (∼0.63 and ∼0.37, respectively). In Aizoaceae, Alooideae and Crassulaceae, modal models fit best in which speciation rate is highest at aridity indices of ∼0.12, ∼0.33 and ∼0.33, respectively, with speciation rate decreasing away from this value. Models with the drift parameter (directionality in character evolution) were best supported in all clades except Agaveae and Alooideae, and were negative. Median aridity index was significantly lower for all seven clades, than for similar-sized clades in the mega-tree of Smith and Brown (2018) (Table 3).

**Figure 4:**
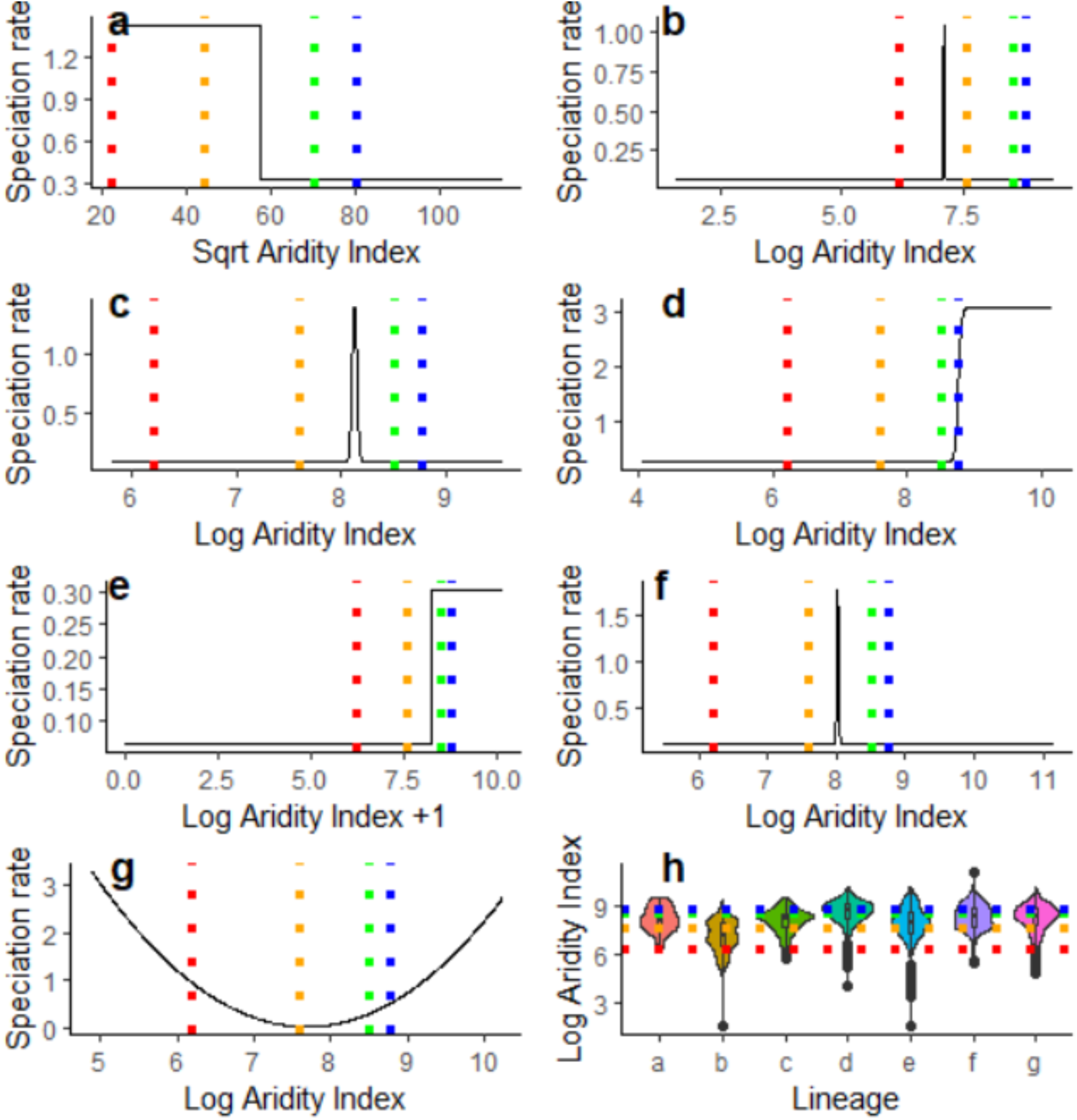
Divergent relationships between Aridity Index and diversification rates in the seven richest succulent lineages. Aridity-dependent speciation is strongly preferred in each lineage, and the best-supported QuaSSE model is plotted (a = Agaveae, b = Aizoaceae, c = Alooideae, d = Asclepiadoideae, e = Cactaceae, f = Crassulaceae, g = *Euphorbia*). Vertical lines indicate UNEP recognised thresholds of Aridity Index (below red = hyper-arid, between red and orange = arid, between orange and green = semi-arid, between green and blue = dry sub-humid, above blue = humid). The distribution of median Aridity Index of species in each lineage (h) is plotted as a violin plot, with mean and boxplot indicated.

**Table 2:** Evidence for aridity-dependent diversification. Best-fitting aridity-index dependent QuaSSE models for each lineage are described, with ΔAIC compared with the null model, and estimated diversification parameters.

| Lineage | Best-fitting model (lowest AIC) | $\Delta AIC$ | Lambda0 | Lambda1 | Midpoint (mode or inflection) | Variance (for modal) | Rate of sigmoidal change | Drift |
| --- | --- | --- | --- | --- | --- | --- | --- | --- |
| Agaveae | Sigmoidal without drift | 36.276 | 1.427596e+00 | 3.282470e-01 | 5.749498e+01 | 8.090703e-01 | 3.186480e+03 |  |
| Aizoaceae | Modal with drift | 214.14 | 6.436245e-02 | 1.076531e+00 | 7.087292e+00 | 1.700798e-04 |  | -0.02517341 |
| Alooideae | Modal | 83.31 |  |  |  |  |  |  |
| Asclepiadoidea<br>e | Sigmoid<br>al with<br>drift | 225.6<br>0 | 0.226763039 | 3.101946406 | 8.754345148 | 40.98085513<br>5 |  |  |
| Cactaceae | Sigmoid<br>al with<br>drift | 168.4<br>5 | 6.431413e-02 | 3.044125e-01 | 8.211629e+0<br>0 |  |  | -5.481608e-02 |
| Crassulaceae | Modal<br>with drift | 173.6<br>0 | 0.107596184<br>7 | 1.813857629<br>1 | 8.016398777<br>4 | 0.000196872<br>9 |  | -<br>0.015317140<br>8 |
| *Euphorbia* | Modal<br>with drift | 147.8<br>1 | 2.187429e+0<br>7 | 2.509818e-02 | 7.684279e+0<br>0 | 2.625149e+0<br>7 |  | -1.574491e-02 |

**Table 3:** T.tests demonstrating that succulent lineages inhabit significantly more arid locations than randomly-drawn similarly-sized angiosperm lineages.

| **Succulent lineage** | **Number of random samples** | **Median aridity index of succulent lineage** | **Mean aridity index of random samples, 95% CI** | **P value** |
| --- | --- | --- | --- | --- |
| Agaveae | 482 | 0.34 | 0.81 (0.73-0.86) | <0.00001 |
| Aizoaceae | 133 | 0.13 | 0.79 (0.73-0.86) | <0.00001 |
| Alooideae | 212 | 0.37 | 0.77 (0.72-0.82) | <0.00001 |
| Asclepiadoideae | 77 | 0.60 | 0.78 (0.71-0.84) | <0.00001 |
| Cactaceae | 52 | 0.30 | 0.75 (0.64-0.86_ | <0.00001 |
| Crassulaceae | 61 | 0.41 | 0.73 (0.66-0.79) | <0.00001 |
| *Euphorbia* | 74 | 0.43 | 0.70 (0.61-0.78) | <0.00001 |

## Discussion

The succulent syndrome has evolved numerous times across angiosperms with different macroevolutionary consequences, from the monotypic family Halophytaceae (Anton et al., 2014) to the megadiverse genus Euphorbia (>2,000 sps) (Nyffeler and Eggli, 2010). Given the highly convergent and environment-dependent nature of plant succulence, it has been tempting to hypothesise that the evolution of the richest succulent clades share patterns as a consequence of coevolution, despite disparate distributions and varied morphologies (Bobich and North, 2009; Alvarado-Cárdenas et al., 2013; Arakaki et al., 2013). These hypotheses have been difficult to test for in the absence of taxonomically-comprehensive time-calibrated phylogenies, as well as reliable datasets on distributions and climatic niches. In this study, we reconstructed well-sampled phylogenetic frameworks for the seven richest succulent clades, time-calibrated with a robust and homogenous approach, and analysed the timings of transitions to succulence, and drivers of diversification. These reconstructions and analyses test the most fundamental hypotheses in succulent plant macroevolution, providing insight into the complex evolution of this strategy in arid biomes, some of the most extensive on Earth and home to remarkable and threatened succulent plants (Pillet et al., 2022).

### Do rich succulent lineage radiations follow a key innovation model?

Under a classic key-innovation model, rapid diversification is triggered upon evolution of some adaptive trait which elevates speciation rate, decreases extinction rate, or both (Slowinski and Guyer, 1993). Despite a general trend for elevated diversification of succulents (Figure 3), our results suggest that succulence might not adhere to a radiation under key-innovation model (Table 1). Accordingly, the phylogenetic distribution of transitions to succulence and rate shifts provides evidence for more nuanced models (Figure 2). Although succulents tend to have faster diversification rates, our results indicate transitions to succulence are generally decoupled from shifts to faster diversification rates, with divergent patterns among lineages (Figure 1 and 2). In addition, there’s also rate shifts distributed in non-succulent branches of five lineages (Agaveae, Aizoaceae, Asclepiadoideae, Crassulaceae and *Euphorbia*). The BiSSE models we employed considered only simple null models of asymmetric diversification (Maddison et al., 2007). We preferred BiSSE models instead of more nuanced HiSSE models (Beaulieu and O’Meara, 2016), because we were simply testing for an association, making no assumption of causality or powerful explanatory power of succulence. While richness is generally elevated by succulence, the mechanism by which succulent radiations are triggered is likely multifactorial.

Our estimated crown dates indicate that the origins of the succulent lineages studied far predate the origins of the arid biomes they inhabit (Table S2). In all cases, phylogenetic analyses have shown that succulent taxa diversify from non-succulent ancestors, leading to the hypotheses that those ancestors were pre-adapted to semi-arid or dry sub-humid micro climates (Hernández-Hernández et al, 2023 *companying paper in review JXB*). The dates we estimated support the interesting hypothesis for the origin of the CAM and C4 carbon concentrating strategies stating that early stem lineages that would give rise to highly specialized groups evolved as a strategy to deal with CO_2_ scarcity, and later colonized and diversified in semi-arid regions (Keeley and Rundel, 2003; Edwards and Ogburn, 2012). In many cases, our reconstructions detect rate shifts occurring more recently than the preceding transition to succulence, however our sampling might be altering the first origin of xeric-adapted taxa within groups. A lag between the evolution of succulence and subsequent radiations is also recovered by previous phylogenetic reconstructions of Cactaceae and Alooideae (Arakaki et al., 2011, Boucher et al., 2020). Recently, the simplicity underlying key innovations and radiations has been questioned and more nuanced models described, in which sequential events are required to elevate diversification (Bouchenak-Khelladi et al., 2015; Donoghue and Sanderson, 2015; Givnish, 2015). The patterns we found adhere to the confluence model described by Donoghue and Sanderson (2015), in which sequential emergence of favourable traits and environmental conditions are required to trigger a radiation.

### The presence of succulence and its effects in diversification rates

The existence of species-poor succulent lineages demonstrates that succulence is not a universal trigger of radiation or diversification rate increases (Olson and Carlquist, 2008; Anton et al., 2014). However, there is a clear and intuitive link in some lineages, with phylogenetic imbalances easily attributable to succulence (Figure 2, 3), and some of these succulent lineages have been identified as belonging to core shifts in angiosperm diversification (e.g. Cactaceae or Aizoaceae, see Magallón et al., 2018). In Cactaceae and Alooideae, simple clade comparisons reveal much higher richness within succulent than non-succulent sister lineages. In the lineages with multiple transitions to succulence (Agaveae, Aizoaceae, Asclepiadoideae, Crassulaceae), BiSSE models support faster net-diversification of succulents (excluding *Euphorbia*). With the exception of *Euphorbia*, we found evidence that succulents diversify more rapidly in all lineages we investigated (Table 1). In depth studies of *Euphorbia* have shown that carbon concentrating photosynthetic mechanisms (CCM) such as CAM or C4 metabolism (determined using carbon isotope ratios) evolved several independent times in the genus, in concordance with increases in diversification rates (Horn, et al. 2014). Since our analyses are focused on a broader characterization based on general morphological traits leading to succulent vs non-succulent, we might be missing the fine details about the physiological strategies present in many non-succulent species included in our seven lineages, obscuring in our study environment-phenotype relationships through evolution in *Euphorbia*.

An association between the evolution of succulence and elevated richness might be intuitive, with succulent adaptations permitting lineages to thrive and diversify in water-scarce environments (Horn et al, 2014; Griffiths and Males, 2017), however in-depth exploration of the components of diversification, speciation and extinction, might reveal a more complex scenario. The elevated net diversification rates of succulents can be shaped by accelerated speciation, decelerated extinction, or both (Donoghue and Sanderson, 2015). Determining which combination drives the rise of succulent lineages is important for our understanding of how the evolution of succulence facilitates species formation. We performed further analyses of the diversification dynamics of the succulent lineages to better understand the mechanisms driving those increases in diversification rates. The results reveal another layer of complexity, with the succulent syndrome showing a different relationship with speciation or extinction rates in different lineages.

For instance, in Agaveae and Asclepiadoideae, elevated succulent net-diversification might be shaped by faster speciation of succulents than non-succulents, with relatively low and similar extinction rates for both states (Table 1). Conversely, succulent speciation rates in Aizoaceae are lower than for non-succulents but succulent extinction is negligible. Crassulaceae supports a more complex model, with faster extinction rate for succulents than non-succulents, but disproportionately faster speciation (Table 1). Our results indicate that the succulent syndrome might be having a different impact on diversification rates in each lineage, contingent on the different environmental contexts and diverging ecological mechanisms such as temperatures (Zhou and Wang, 2016), precipitation (Li et al., 2016), elevation, soils and floristic competition (Abd El-Ghani, 2008; Talmon et al., 2011). Furthermore, locally varying factors are known to shape patterns of Cactus endemism (Trabuco Aramal et al., 2022), extinction rate (Pillet et al., 2022) and speciation rate (Thompson et al., 2023), a pattern is likely to extrapolate across succulent lineages.

### Diverse macroevolutionary responses to aridification and CO_2_ concentrations

Perhaps the most fundamental hypotheses we were able to test in this study is for a relationship amongst the diversification of succulents and aridity. In the absence of a fossil record or reliable reconstructions of global aridification processes during the Earth history, we relied on a comprehensive phylogenetic approach and current aridity index values (Trabucco and Zomer, 2018). Although a dependence of diversification on aridity is clearly supported by our results (Table 2), the nature of the relationship varies remarkably among lineages (Figure 4). The level of aridity in which speciation rates are most rapid ranges from hyper arid in Agaveae, Alooideae and some *Euphorbias*, to more humid in Asclepiadoideae, Cactaceae and other *Euphorbias* (Table 2, Figure 4). Arid and semi-arid regions are the most extensively distributed over the Earth surface (Perry et al, 1979), and thus, arid adapted biotas might be the most abundant and widely distributed. It is not surprising that most of the species included in our study inhabit them, in comparison with a random sample (Table 3). In particular, mean aridity index values for succulents correspond to semi-arid biomes, given their clear tendencies towards environments with less water availability (Table 3). Average AI values however, might not reflect the versatility of the succulent syndrome, since it’s been shown that within the same lineage, succulents might inhabit a wide range of climatic niches (Hernández-Hernández et al, 2023 *companying paper in review JXB*). The lack of a linear relationship among the dynamics of diversification of succulent lineages and AI suggests that different succulent lineages respond differently to this climatic variable (Figure 4). This lack or weakness of a linear relationship among aridity variables and diversification rates in succulents has previously been found in macroevolutionary studies using a different statistical approach (PGLS, see Hernández-Hernández et al, 2023, *companying paper in review JXB*). These results might be puzzling under an assumption that an elevated WUE through water storage is an adaptation towards surviving arid environments.

Plants that evolved the succulent CAM strategy are not only highly efficient in the use of water. This type of photosynthesis comes up with a high efficiency in photosynthesis by enhancing carbon fixation through mechanisms that concentrate it around Rubisco, suppressing photorespiration (Borland et al., 2014). As a specialized CCM, the CAM strategy has been suggested to evolve in response to decreases in environmental CO_2_ concentrations (Arakaki et al. 2011), in particular in response to the collapse of atmospheric CO_2_ commencing ∼15 Mya, and the global expansion of arid environments in the Late Miocene (Tripati et al., 2009). The relation among CO_2_ drop-off in concentrations and plant radiations was confirmed previously for CCM Euphorbias, both CAM or C4 (Horn et al, 2014) and also widely documented in C4 grasses (Spriggs et al, 2014; Bellasio et al, 2022).

In addition to observing the distribution of the estimated times of rate shifts and times of appearance of succulence, we investigated the impacts of CO_2_ with different, but complementary approaches. Similarly as with aridity values, we found mixed support for dependence of lineage-wide dynamics on atmospheric CO_2_. Only Asclepiadoideae and *Euphorbia* are strongly supported as dependent on continuously-varying paleoclimatic CO_2_, while the remaining lineages support either time-dependence, or neither time nor CO_2_ (see Results section). Even more conflict is found in investigations of episodic diversification on either side of the hypothesised collapse in CO_2_ ∼15 Mya. Again, only Asclepiadoideae and *Euphorbia* demonstrate differential tree-wide diversification on either side of ∼15 Mya, with rates for *Euphorbia* flouting the expectation of higher diversification rates in recent time, confirming previous results (Horn et al, 2014). However, uncertainty in estimated rates following ∼15 Mya is wide and overlaps with the distribution of earlier rates. Evidence for CO_2_-dependent diversification of succulents is at best unequivocal, with lineage-wide variation generally uncoupled from CO_2_ with the exception of Asclepiadoideae. Furthermore, the uncertainty surrounding divergences of succulent clades renders any analyses which require accurate time-calibrations difficult. The reconstructions we presented were calibrated by robust estimates from a comprehensive analysis of angiosperm divergences (Ramírez-Barahona et al., 2020), but we were not able to account for uncertainty, which is wide in many estimates.

### The ecological meaning of the succulent syndrome

We believe that the macroevolutionary patterns we observe evidence both the ecological as well as environmental complexity of the semi-arid and dry-subhumid regions where succulent lineages diversified, and suggest that the idea of convergence towards similar climatic conditions need to be reassessed or better specified (Alvarado-Cárdenas et al, 2013). Climatologists and macroecologists have long agreed that, although the general landscape and the functional composition of floras might be similar, different arid and semi-arid regions of the world are climatically very different (Walter, 1973; Perry et al, 1979; Ellenberg, 1981). These ideas are supported by our estimated dates, as well as dates previously reported, indicating that succulents originated in time frames that predate aridification onsets, diversifying later in environments already semi-arid (Horn et al, 2014; Hernández-Hernández et al, 2014; Hernández-Hernández et al, 2023 *companying paper in review JXB*).

In addition to the complex climatic context for the evolution of succulents, the macroevolutionary patterns we observe might also evidence our lack of understanding of the ecological complexity of the succulent strategy beyond survival in water scarcity. Some previous results of studies with specific lineages show that the origin and radiation of succulents might be driven by more complex ecological fosters than aridity per se. For example, growth form and pollination syndrome might have favoured speciation in several cacti lineages (Hernández-Hernández et al., 2014); the high lability of life-form in *Crassula* might have allowed the colonization of new niches (Lu et al., 2022); whereas edaphic and topographic complexity might have favoured increases in speciation rates in Aizoaceae (Ellis et al., 2006; Valente et al., 2013). A recent study of climatic niches of succulents versus non succulent relatives within the Order Caryophyllales confirmed the versatility of the succulent strategy, with a weak relationship of diversification rates and aridity, and estimated dates showing that the origin of succulents predates the aridification of the biotas under which they diversified (Hernández-Hernández et al. *companying paper in review JXB*). These data suggest a broader and unexplored ecological relevance of the succulent strategy. For example, it has been suggested that the evolution of succulence might provide these lineages the possibility to explore the reproductive niche space, and alter phenologies to reproduce during the dry season (Herrera, 2009; Hernández-Hernández et al. *in review JXB*).

## Conclusions

Although there is an emerging interest in the functional mechanisms underlying the physiological and metabolic components underlaying the CAM photosynthesis – a key component of the succulent syndrome – (e.g., Heyduk et al, 2022; Winter and Smith, 2022; Li et al, 2023), questions remain about the macroevolutionary trajectories leading to the succulent syndrome, or its ecological significance. For example, we have no information about when succulence arose and the importance of aridity and climate in diversification remain unresolved (Grisebach 1872, Ellenberg, 1981; Von Willert et al; 1992). In general, our results show a lack of concordance among macroevolutionary patterns underlying convergent evolution of succulence among these clades. Generally, richness is elevated in succulent lineages compared to non-succulent sister lineages, but state-dependent diversification is not supported in every case, and most rate-shifts are decoupled from transitions. We found no synchronicity in the origin and diversification of succulent lineages, and a lack of support for hypothesis of succulents as global organismal responses to general aridification trends (Arakaki et al, 2011). In general, shifts in diversification rates and shifts towards succulence occurred mostly in the last 15Mya (Miocene), however, a recent general increase in diversification rate might be ubiquitous to all angiosperms (Magallon et al, 2018). As puzzling results, diversification rate increases, shifts towards succulence and/or increases in species numbers do not necessarily co-occur, indicating that the ‘succulent key innovation’ should be reassessed or specified and that it might be contingent on diversity of environmental challenges or ecological interactions.

## Acknowledgements

JBT would like to acknowledge Roger and Sue Whorrod for providing a research scholarship, a Santander travel grant for facilitating collaborative research with UNAM, and Bjorn Kopperud for providing advice with a TESS model. Further acknowledgments go to GMK for comments and suggestions to preliminary versions of this manuscript.

